# scExploreR: a flexible platform for democratized analysis of multimodal single-cell data by non-programmers

**DOI:** 10.1101/2025.05.28.656649

**Authors:** William M. Showers, Jairav Desai, Stephanie R. Gipson, Krysta L. Engel, Clayton Smith, Craig T. Jordan, Austin E. Gillen

## Abstract

Single-cell sequencing has revolutionized biomedical research by uncovering cellular heterogeneity in disease mechanisms, with significant potential for advancing personalized medicine. However, participation in single-cell data analysis is limited by the programming experience required to access data. Several existing browsers allow the interrogation of single-cell data through a point-and-click interface accessible to non-programmers, but many of these browsers are limited in the depth of analysis that can be performed, or the flexibility of input data formats accepted. Thus, programming experience is still required for comprehensive data analysis. We developed scExploreR to address these limitations and extend the range of analysis tasks that can be performed by non-programmers. scExploreR is implemented as a packaged R Shiny app that can be run locally or easily deployed for multiple users on a server. scExploreR offers extensive customization options for plots, allowing users to generate publication quality figures. Leveraging our SCUBA package, scExploreR seamlessly handles multimodal data, providing identical plotting capabilities regardless of input format. By empowering researchers to directly explore and analyze single-cell data, scExploreR bridges communication gaps between biological and computational scientists, streamlining insight generation.

## Introduction

Single-cell sequencing has provided transformative insights into cellular heterogeneity, contributing to breakthroughs in the fields of cancer (1–3), cardiovascular disease (4, 5), and psychiatric conditions (6, 7). Single-cell sequencing shows great promise in developing new therapeutics and realizing personalized medicine, but interpretation of data is difficult and requires deep technical expertise. Programming experience in either Python or R is required to access single-cell data, often necessitating collaborations between researchers and bioinformaticians. There is some overlap in biological expertise between researchers and bioinformaticians, but differences in perspective between the two fields lead to barriers in communication, complicating the process of insight generation from these rich datasets.

Several single-cell visualization tools have been developed, but many tools are limited in terms of ease of use, flexibility of input data, plotting capabilities, or the ability to perform differential gene expression (DGE) analysis. For example, the widely used UCSC Cell Browser (8) only produces dimensional reduction plots, feature plots and violin plots, and does not allow users to “split” plots into groups of cells to make comparisons. ShinyCell (9) produces many visualization types used in single-cell analysis, but can’t perform differential gene expression. Loupe (10) can perform differential gene expression and excels in both the depth of customization of plots and the ability to subset datasets, but only supports the output format of the 10X Genomics CellRanger pipeline, or Seurat objects via conversion.

To combine the strengths of existing single-cell visualization tools, we developed scExploreR. scExploreR is an analysis platform built on R Shiny (11) that enables in-depth visualization and differential expression analysis of single-cell data in a graphical user interface (GUI). The programming barrier to single-cell data analysis is removed with scExploreR, and researchers are able to participate directly in common single-cell analysis tasks. Communication between researchers and bioinformaticians still occurs, but is focused on advanced analysis tasks, as a follow-up to findings from using scExploreR. Bioinformaticans also remain involved in the processing of single-cell data and deployment of the platform. scExploreR’s interface is intuitive, with tooltips and extensive documentation for features. Visualization tasks are facilitated by a modular interface design, with consistent menu elements across plot types. The interface for differential expression analysis is simple and clearly labeled, facilitating the selection of cell populations for comparison. scExploreR is flexible in terms of the classes of input objects, supporting Seurat (12), SingleCellExperiment (13), and anndata (14) classes. Multimodal data is explicitly supported by scExploreR. Any single-cell modality that can be expressed as a matrix of counts per cell, per gene, and included in a single-cell object, can be loaded in scExploreR. These features allow users to perform in-depth single-cell analysis without requiring programming expertise, making scExploreR an accessible and powerful tool for researchers.

## Materials and Methods

### Setup and Deployment

scExploreR is a Shiny (11) app written in R and distributed as an R package. The software is freely available from our GitHub repository (15). Formatting the app as a package allows for quick deployment: the app is run simply by loading the package and executing an R function. Instructions for installing the package and its dependencies are available in the README file in the repository, and more detailed instructions are available on our website (16). Using the app requires no programming experience, but some programming is required to configure and deploy the app. We envision bioinformaticians with programming expertise and familiarity with the dataset taking responsibility for setting up scExploreR instances. We define this type of user as an “admin” in this manuscript, and we refer to users interacting with the dataset via scExploreR as “end users”. scExploreR may be deployed in several different contexts, for different purposes. The app may be deployed locally to visualize a dataset for quick exploration, or for an informal presentation to colleagues, or it may be deployed on a server to share single-cell data with a group of colleagues. Admins may also use Docker (17, 18) to containerize an app instance with their data to streamline deployment on a server. The containerized app instance can then be distributed to collaborators to run the app and view data. Since Docker containers function the same on all operating systems, any collaborator who downloads and runs the container will be able to view the dataset in the scExploreR instance. We provide instructions for creating a docker container on our website (19). If the app is deployed on a server, admins are responsible for configuring access controls and security settings.

### Input Data

scExploreR supports single-cell datasets saved in the Seurat (12), SingleCellExperiment (13), or anndata (14) object classes. Support for these object classes is achieved via our SCUBA (20) R package. Because of the use of SCUBA, visualization capabilities in scExploreR are identical across object types. Data from any single-cell omics modality that can be expressed as a counts matrix and stored in these object classes can be loaded in scExploreR. The app is designed to work with an object that has been through a pre-processing workflow (dimensional reduction, clustering, cell type annotation). Objects that have not undergone these steps may be loaded in the browser as long as they’re in the previously mentioned formats, but functionality may be limited without the information computed during pre-processing.

Single-cell objects in Python are loaded using Reticulate (21). To load Python single-cell object classes in scExploreR, admins must have a Python installation with the following packages installed: Numpy, Pandas, Scipy, Anndata, and Scanpy. We recommend using anaconda to manage Python installations, and provide instructions for doing so in our GitHub README (15).

### Configuring Objects

Before loading single-cell objects into an instance of scExploreR, admins must first configure object-specific settings via an additional Shiny app provided with the scExploreR package. Admins first run the config app via the function `run_config`, using the path to the object as input (**Figure 1A**). The config app has several categories of settings organized into tabs. The general tab is used to set the display name of the objects when hosting multiple objects on the server, to provide a description of the dataset for end users, and a preview dimensional reduction plot to show on the datasets screen of the main app. The “Assays” and “Metadata” tabs are used to choose the modalities and metadata variables to display to end users, respectively, and to define settings specific to each. Settings defined in the config app are downloaded in a YAML file, which is supplied with the object when starting up an scExploreR instance.

**Figure 1.**
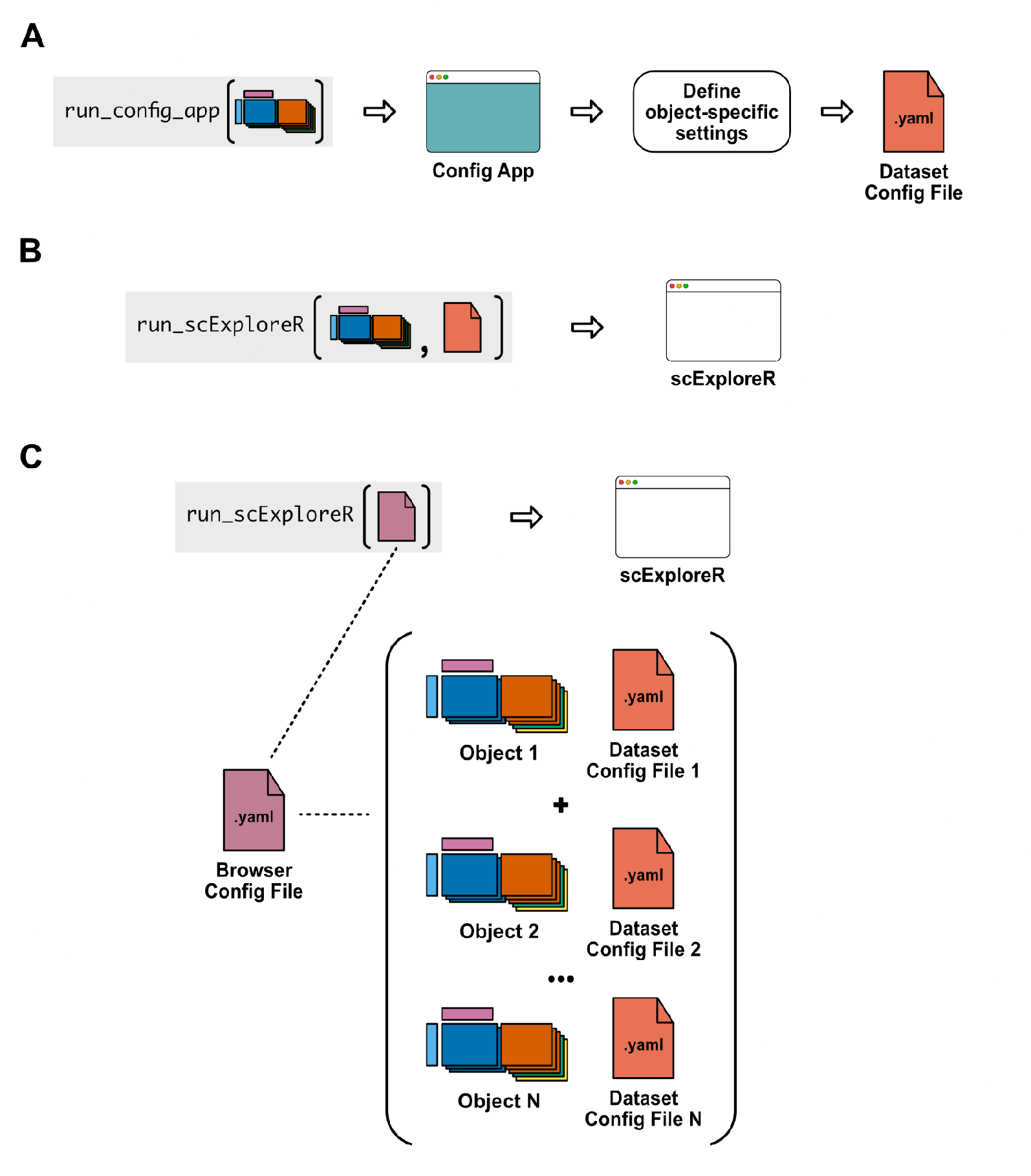
scExploreR is easy to deploy and can be set up with one or many single-cell objects. (**A**) To load an object into scExploreR, users start by running the config app on the object to define object-specific settings. The config app is run by calling a single R function, `run_config`. The config app saves settings to a .yaml config file. (**B**) To deploy scExploreR with a single object, users simply call `run_scExploreR` with paths to the object and config file. (**C**) To deploy an instance with multiple objects, users may write a “browser config” .yaml file defining the paths to each object and config file. Users then call `run_scExploreR`, passing just the browser config file as input.

### Initializing an App Instance

After defining the settings for each object in the config app, admins run an instance of the main app using `run_scExploreR`. To initialize the app with a single object, the paths to the object and the config file are supplied directly to the function (**Figure 1B**). For multi-object deployments, an additional YAML file is created defining the paths to each object and config file (**Figure 1C**), as well as settings specific to that deployment. This “browser config file” is then passed to the `run_scExploreR` function.

## Results

### Comparison with Existing Single-Cell Visualization Platforms

**Figure 2 A-D** compares scExploreR with eight single-cell visualization tools: UCSC Cell Browser (8), ShinyCell (9), Loupe (10), CELLxGENE (22), Cirrocumulus (23), Vitessce (24), iSEE (25), and Kana (26). Among these, scExploreR stands out for its versatility in both visualization and DGE analysis (**Figure 2A**). Of the platforms that can perform DGE, scExploreR offers the greatest versatility in visualization, and is one of few tools with explicit support for multimodal data. Any modality that can be expressed as a matrix of counts per cell, per gene, and included in a single-cell object, can be loaded in scExploreR. In addition to the variety of single-cell modalities that can be analyzed and the variety of plot types that can be produced, scExploreR allows for extensive customization of plots (**Figure 2B**). scExploreR is one of few tools that allows for control over both high-level aspects of plotting, such as “splitting” plots by a metadata variable, and more detailed aesthetic aspects, such as color palettes and setting the size and resolution of image downloads. scExploreR is flexible in terms of the classes of input objects, supporting Seurat (12), SingleCellExperiment (13), and anndata (14) classes (**Figure 2C**). All supported classes are natively supported in scExploreR as opposed to providing a conversion function to an accepted format. scExploreR is also flexible in terms of the context in which it may be deployed (**Figure 2D**). scExploreR can be deployed locally or on a server with any number of datasets, and it is one of few tools that can be deployed with Docker.

**Figure 2.**
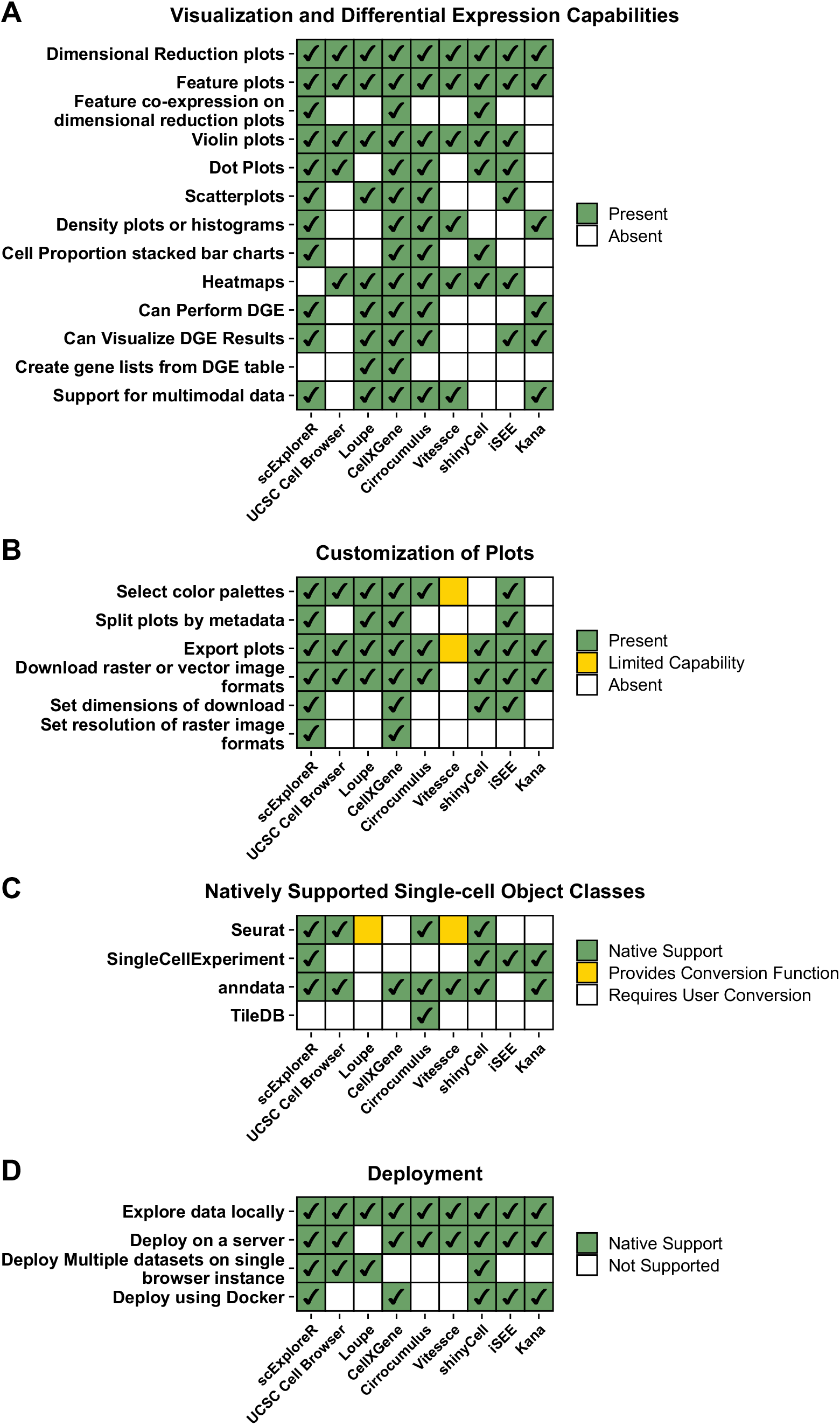
Comparison of scExploreR with commonly used single-cell browsers. Single-cell browsers were compared on the criteria indicated in A-D. Features in each category are indicated as being present, absent, or present in a limited capacity, based on the colors in the legend. (**A**) Of all browsers that support differential gene expression, scExploreR is the most versatile in plotting capabilities. scExploreR also explicitly supports multimodal single-cell data. Data from any single-cell modality can be used as input for scExploreR if it can be expressed as a cell by feature matrix. (**B**) scExploreR allows for creation of publication quality plots from single-cell data. scExploreR offers extensive choices for customizing and exporting plots. Plots can be exported in vector or raster image formats, and the image dimensions and resolution can be set. (**C**) scExploreR was designed to natively support many commonly used single-cell object classes. Native support, as opposed to providing a conversion function to a format used by the browser, minimizes the risk of data loss. (**D**) scExploreR can be deployed in a variety of contexts. Instances of scExploreR can be deployed in Docker containers to manage cross-platform deployment.

### Visualizations

The interface of scExploreR is split between three tabs: the Info tab, the Plots tab and the Differential Expression tab. The Info tab displays general information, dataset specific information, and dataset controls. In the Plots tab, end users may produce and export several plot types commonly used in single-cell analysis. Plots are produced in R via ggplot2, and many plot types are based on the MIT-licensed open-source Seurat (12) package (with attribution). The plot types that can be produced by scExploreR are shown in **Figure 3**. To summarize a dataset, scExploreR produces dimensional reduction plots (DimPlots), and cell type proportion plots. DimPlots color cells by categorical metadata like clusters or sample names on a two-dimensional reduction (UMAP, PCA, etc.).. Cell type proportion plots show cluster proportions by a sample or patient level variable, and are useful for comparing proportions across samples or patients. To visualize and compare feature expression profiles, scExploreR produces feature plots, violin plots, ridge plots, and dot plots. The variety of plots allows end users to visualize feature expression in different ways. Feature plots show high-level trends in expression, while violin and ridge plots visualize the distribution of expression in more detail. Dot plots provide additional detail, visualizing average expression within defined groups of cells, as well as the percentage of cells within each group with non-zero expression of a feature. Dot plots are also particularly well suited to the visualization of large numbers of features at once. scExploreR also produces scatter plots, which are useful for visualizing correlation between two features. All plots support multimodal data. Features may be genes, surface protein markers, chromatin accessibility measurements, or any other form of single-cell data that can be expressed as a per-cell counts matrix. End users enter features for potting in a search interface, where features are labeled with the name of the associated assay/modality, as defined in the config app.

**Figure 3.**
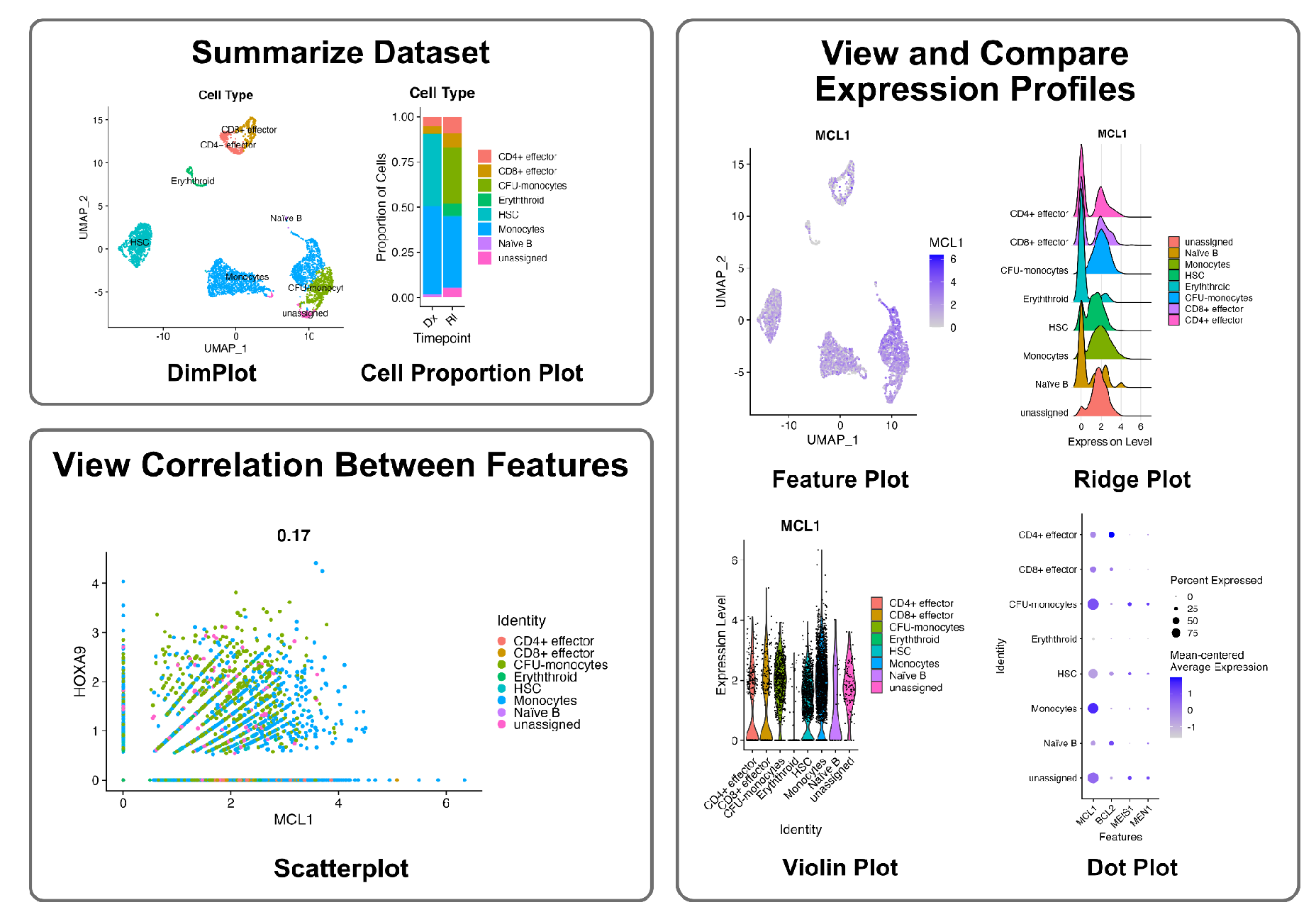
scExploreR produces visualizations commonly used in single-cell data analysis. The plot types produced by scExploreR are shown and grouped by function. To summarize datasets, scExploreR produces dimensional reduction plots (DimPlots) and Cell Proportion plots. DimPlots provide a general summary of the data, while cell type proportion plots are useful for a per-sample summary. Several plots are provided to examine trends in feature expression. The term “feature” may refer to genes, surface proteins, or other modalities. Feature plots show a high-level overview of feature expression by cell type, while ridge and violin plots show the distribution of expression, by groups of cells chosen by the user. Dot plots are useful for large numbers of features and show the average expression per cell group, as well as the percentage of cells with non-zero expression of the feature. Correlations between features may also be analyzed using scatterplots.

In addition to the variety of plot types, scExploreR offers extensive options for customizing each plot type to create and download publication quality figures. A schematic of the Plots tab interface along with some operations that may be performed is shown in **Figure 4**. Plots in the Plots tab are sorted by type, and enabled by clicking the switch next to a plot type in the top left corner of the window. Options for plots display in the left side of the window, and are also sorted by plot type. Options “panels” are modular in that they use a consistent style across plot types with many elements in common for ease of use. Most plots have a “group by” selection, which allows end users to specify a metadata variable used for grouping cells on the plot, and a “split by” selection, which facets groups of cells into separate panels. For plot types where groups are represented in order in the plot (i.e. violin, ridge, dot, and proportion plots), an interface is provided to sort groups. End users may sort groups in ascending or descending alphabetical order, or by average feature expression (or in the case of cell type proportion plots, the relative proportion of a cell type in the group). For DimPlots and Feature Plots, end users may switch between alternate dimensional reduction projections that are included in the object and enabled in the config app by the admin. End users may also choose the metadata used to label cells. For Feature Plots with two features, end users may either view feature plots for these genes side-by-side, or enable a “co-expression” plot, where each feature is assigned a separate color, and the degree to which both features are expressed in each cell is given by a “blended” color for that cell. Several aesthetic options may be adjusted, such as the plot title, color palettes, the appearance of legends on the plot, or the number of columns in the layout for faceted plots. All plots may be downloaded in either raster (.png) or vector (.svg) formats. The dimensions of downloads may be set in inches, and the image resolution may be set for raster images.

**Figure 4.**
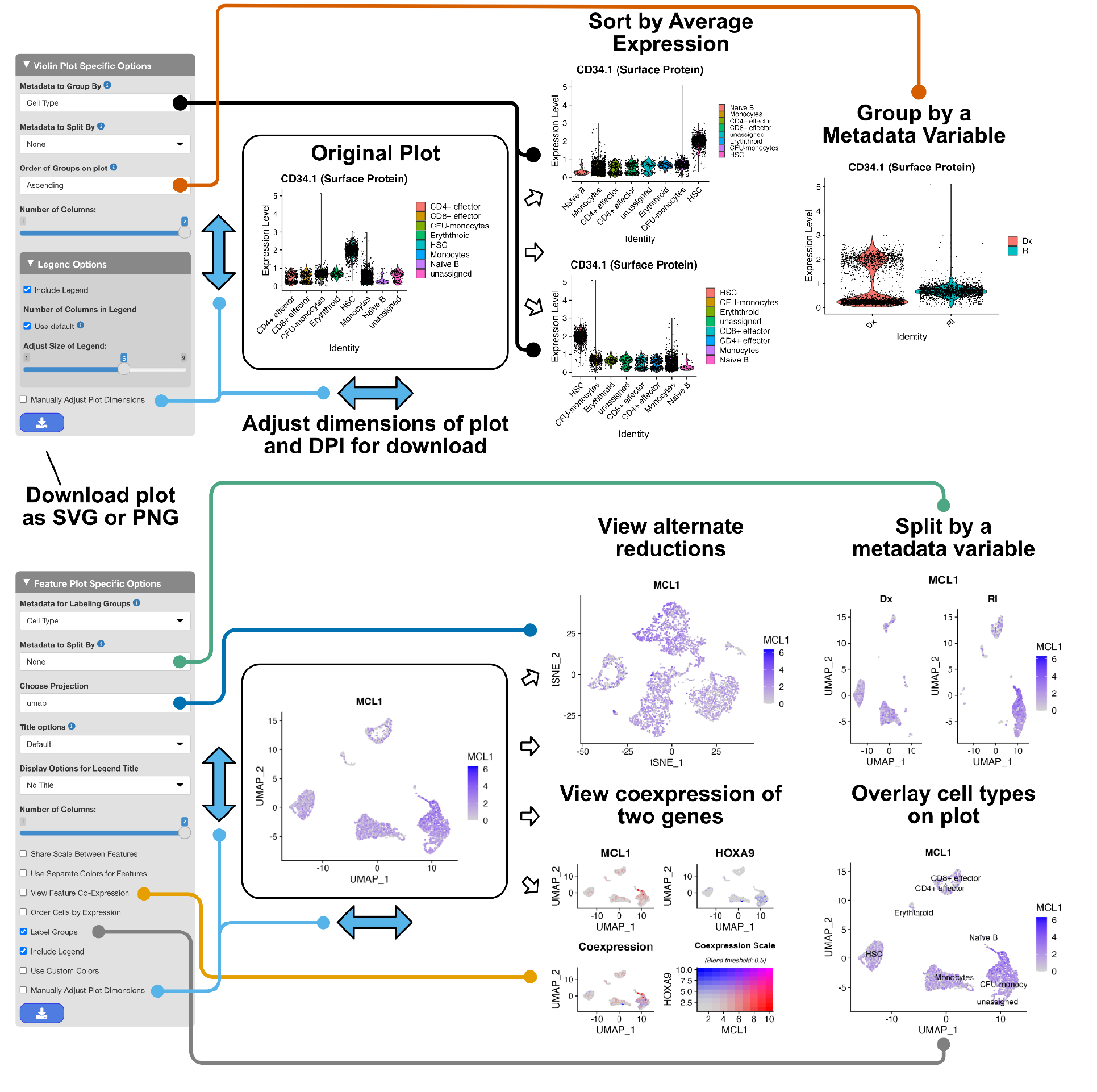
scExploreR provides extensive options for creating and downloading publication-quality figures. A violin plot and feature plot are shown to demonstrate examples of operations that may be performed. Screenshots of the interface for choosing options for each plot are shown on the left hand side. To the right of each interface, default plots are shown along with operations that can be performed and the results of those operations. Connectors link each operation with the corresponding interface menu to perform the operation. Users may choose a metadata variable to define groups of cells on the plot (orange connector), and on many plots users may split/facet plots by a metadata variable (green connector). On plots involving comparisons of feature expression across groups of cells, users may sort groups in ascending or descending order by feature expression (black connector), in ascending or descending alphanumeric order, or according to a custom order. On feature plots and DimPlots, users may select the reduction coordinates to display (blue connector) or metadata to use for labels overlaying the plot (gray connector). For feature plots with two features, users may either view feature plots for these genes side-by-side, or enable a “co-expression” plot (yellow connector). Co-expression plots assign a separate color to each feature, and “blend” these colors together according to the degree to which each feature is expressed in each cell. Users may download plots in either SVG or PNG format, and can set the size of the plot in inches as well as the resolution of the downloaded image (light blue connector).

### Subsetting

By default, scExploreR shows all cells in the single-cell object for visualization and analysis, but end users can define subsets to focus on specific cell types or samples. Subsetting is achieved through a flexible interface that allows criteria to be set based on categorical metadata, numeric metadata, or feature expression. Criteria are set one at a time, with a single criterion consisting of a single metadata variable or feature, and the values for that variable or feature for which cells should be included. For example, one criterion may be cells where “cell type” is equal to “T-cells”, “NK cells”, or “monocytes”, or cells where expression of “CD34” is greater than or equal to a defined threshold. For criteria based on numeric metadata or feature expression, thresholds are defined using an interface with a ridge plot of the feature or metadata variable. Criteria are defined one at a time, and all criteria are joined by AND logic. If end users have a programming background, they may also define criteria with code, by passing an R expression interpretable by the `subset` parameter of the `subset()` method of SeuratObject. Code-based subsetting is currently only supported for Seurat objects.

### Differential Gene Expression Testing

In the Differential Expression tab, end users may conduct differential expression tests and view results in real time. Differential expression tests use a Wilcoxon rank sum (27) method. For Seurat objects, scExploreR uses the presto package (28) to run the test, and for Seurat objects with BPCells matrices, the `marker_features` function from BPCells is used. For anndata objects, testing is performed via Scanpy, using the `scanpy.tl.rank_genes_groups` function. Testing via a Wilcoxon rank sum method was chosen for performance, but in objects with multiple samples and imbalances in the number of cells per patient, a pseudo-bulk approach is more appropriate (29). Differential gene expression testing is not supported for SingleCellExperiment objects.

End users follow a three-step process to identify the comparisons they wish to run in the Differential Expression interface (**Figure 5**). End users first choose the type of comparison to perform, from either differential expression or marker identification. For differential expression, end users identify two groups of cells to compare, and genes that are differentially expressed in one group vs. the other will be returned. For marker identification, end users split cells into three or more groups, and genes that are differentially expressed in each group vs. all other cells are returned. Next, end users define the groups of cells to compare. For marker identification, end users select a categorical metadata variable from the options enabled in the config app, and groups will be created for each value within that variable. For differential expression, end users may define groups using either metadata or feature expression. To define groups by metadata, end users choose a variable to use for forming the groups, and then select the values within that variable to use to define either the first and second group. End users may select a single value for a group (for example, to compare two cell types) or multiple values (to compare a set of samples with a second set of samples). To define groups by feature expression, end users select a feature to use, and define an expression threshold using an interactive ridge plot interface. Cells with expression greater than or equal to the threshold will be placed in one group, and cells with expression less than the threshold will be placed in the other. After defining groups, end users next choose to run the test in the full object, or a subset of cells. The subsetting interface functions as described in the “subsetting” subsection, and groups are created from the cells that match the criteria entered.

**Figure 5.**
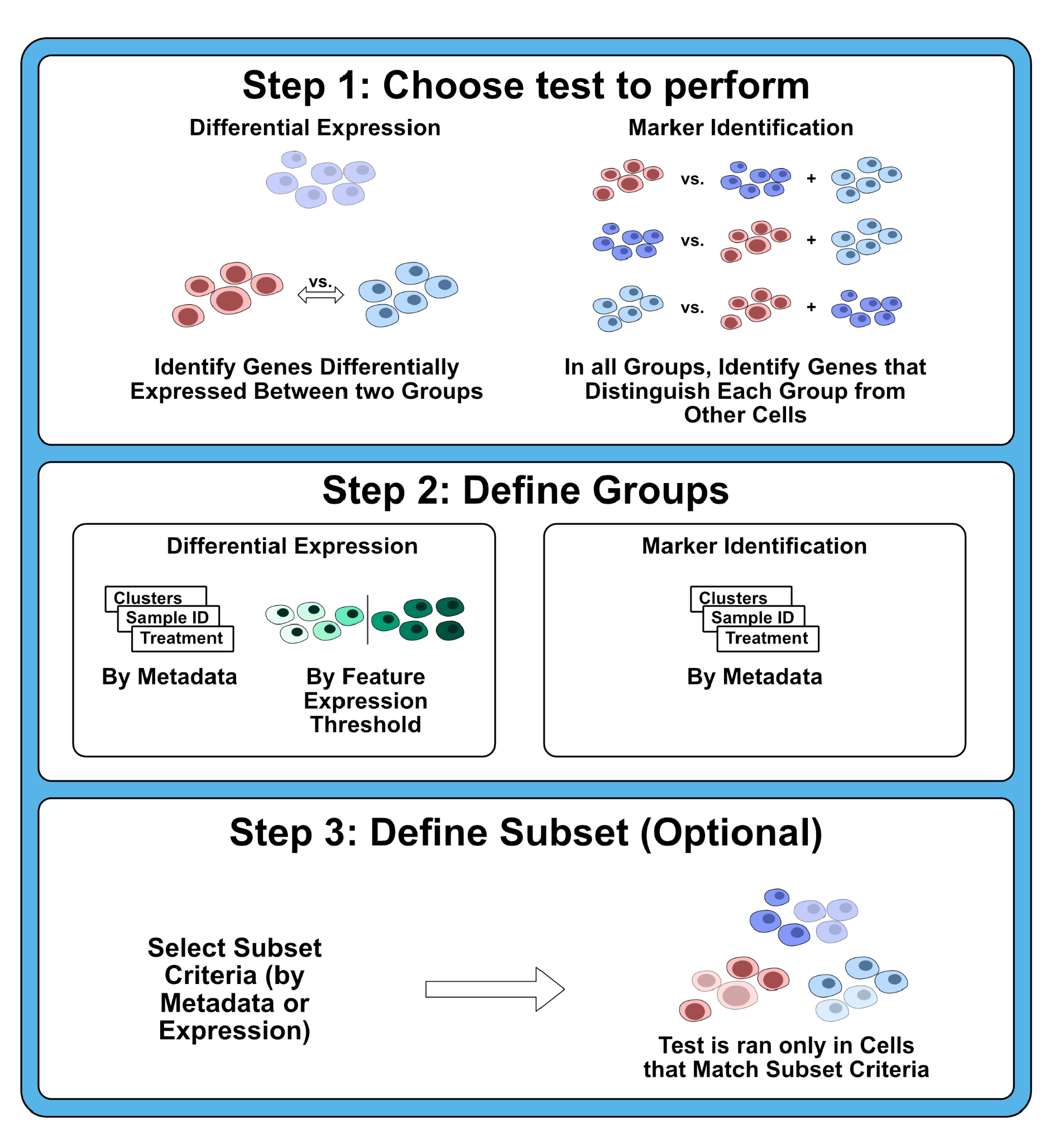
Differential gene expression test selection workflow. The process used to run differential gene expression tests in scExploreR is shown. Users first select the type of test to perform, from either differential expression or marker identification. For differential expression, two groups of cells are compared, and genes differentially expressed in each group are returned. For marker identification, three or more groups are compared, and genes that distinguish each group from all other cells are shown. Next, users define the groups of cells to use for the test. For differential expression, users may define groups based on either metadata or feature expression. To define groups by metadata, users choose a metadata variable to use for the comparison, and then users define two groups using values of that variable. Users may select a single value, or multiple values for either or both groups. For definition by feature expression, users select a feature, and then define a threshold by making a selection on a ridge plot for that feature. Cells with expression less than the chosen threshold will be compared with cells with expression greater than or equal to the threshold. For marker identification, groups are constructed from metadata. Groups are created from each of the values present in the selected metadata variable. Finally, users may optionally select a subset of cells in which to perform the test. Subsets may be created based on feature expression thresholds or categorical metadata.

The interface of the Differential Expression tab is shown in **Figure 6**. On the upper left side of the screen, end users define and run the differential gene expression test. Once a test is complete, summary statistics display on the top of the screen, and the table of differentially expressed genes displays beneath the summary statistics. In the table, one row is shown per gene, per group in the comparison, unless the end user indicates to show only genes that are upregulated in groups. End users may use either the filter text boxes in the table or the filter interface on the left side of the screen to identify genes differentially expressed in each group based on an adjusted p-value cutoff, or they may enter a gene to see if it is differentially expressed in a group, or in which groups the gene is differentially expressed. End users may export the table as a CSV file.

**Figure 6.**
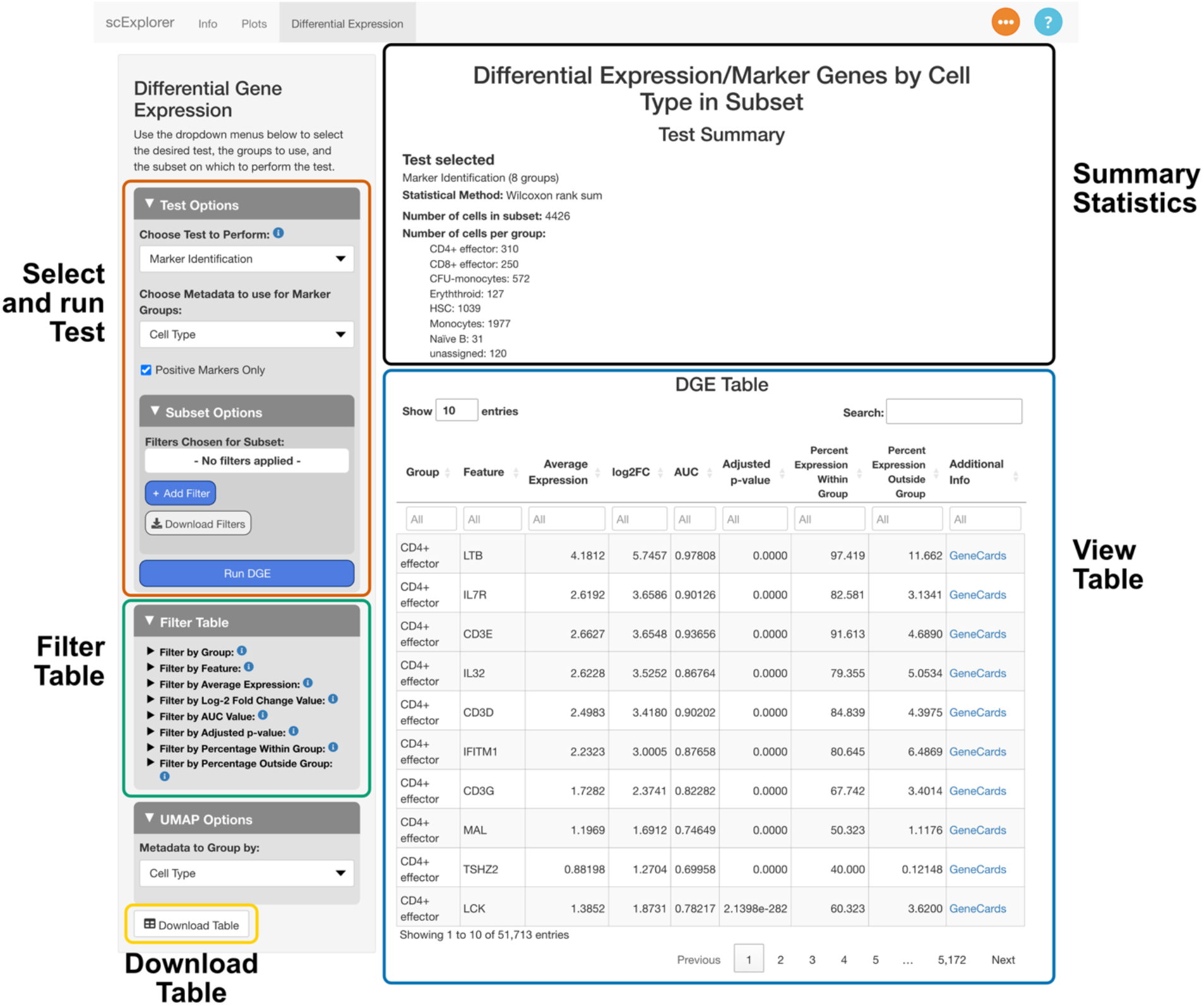
Layout of differential gene expression tab. Users select the desired test in the top left corner (orange outline). Initially, this is the only interface shown in this tab, and other elements are shown when a test is completed. In the main window on the right, users can view a summary of the test performed, the groups chosen, and the number of cells in each group (black outline). Beneath this, the table of markers/differentially expressed genes is shown (blue outline). The table shows results by cell group, along with statistics for each result. The table can be filtered using the interface on the left (green outline), and it can also be downloaded to a .csv file (yellow outline).

## Discussion

scExploreR is a web app focused on the production of visualizations commonly used in single-cell data analysis, and differential gene expression analysis. scExploreR stands out from similar tools in the depth of analysis that can be performed without code, and its ease of use. Visualization capabilities are particularly extensive, allowing end users to produce Seurat-style (12) visualizations without needing to write R code. Common R and Python single-cell object classes may be loaded in scExploreR, and single-cell data from any modality that can be represented as a counts matrix may be loaded into scExploreR. The browser can be deployed with minimal coding experience locally or on a server, and instructions are provided for cross-platform deployment in Docker. scExploreR’s interface is intuitive and consistent across app functions.

Future development will focus on expanding scExploreR’s functionality to address emerging needs in single-cell analysis. Planned features include explicit support for spatial transcriptomics data, which would enable end users to visualize spatially resolved gene expression patterns. For marker identification tasks, we also plan to implement the approach in Scran’s (30) `findMarkers()`, which improves upon the current Wilcoxon rank sum implementation by aggregating p-values across replicates. We also plan to add the ability to automatically transfer genes from the differential expression tab to the plots tab, and the ability to save and load lists of genes and subsets.

We anticipate scExploreR will facilitate interactions between bioinformaticians and non-informatics researchers, with bioinformaticians deploying the app with data they have processed, and non-informatics researchers analyzing deployed data. scExploreR is also of use to bioinformaticians as it dramatically speeds up initial analysis and exploration of a dataset. scExploreR bridges the accessibility gap in single-cell data analysis, making it an invaluable resource for advancing single-cell research. Its intuitive interface, robust visualization tools, and flexibility for multimodal datasets empower researchers to directly engage with their data, fostering new discoveries and accelerating insights. By broadening participation in single-cell data analysis, scExploreR boosts efficiency of research and broadens the potential for insight generation in any field where single-cell data is used.

## Competing Interests

A.E. Gillen and C.T. Jordan report other support from RefinedScience outside the submitted work.C.A. Smith is the Chief Innovation Officer of RefinedScience and the Chief Medical Officer of Oncoverity and receives financial support from both. No funding from Oncoverity was used in this work. No disclosures were reported by the other authors.

The contents do not represent the views of the U.S. Department of Veterans Affairs or the United States Government.

### Grant Information

This work was supported in part by Career Development Award #IK2BX004952-01A1 from the United States (U.S.) Department of Veterans Affairs Biomedical Laboratory Research and Development Service to AEG, US NIH R35CA242376 to CTJ, and Leukemia and Lymphoma Society DGP (8052-25) to CTJ and AEG.

## Acknowledgements

The authors would like to acknowledge Monica Ransom, Sarah E. Staggs, Kent Riemondy, Stephanie Redmond, Devin Burke, Abbigayl Burtis, Kay Linker, Mark J Althoff, Maria Amaya, Shanshan Pei, Ian Shelton, Brett Stevens, Sweta Patel, Ana Vujovich, Anagha Inguva, Mohd Minhajuddin, Maura Gasparetto, and Regan Miller for their thoughtful comments and suggestions during the development of this package and the writing of this manuscript.

## Data Availability

Source code for scExploreR is available from our GitHub repository at https://github.com/amc-heme/scExploreR.

## Author Contributions

Conceptualization: BS, AEG; Funding acquisition: AEG, CS, CTJ; Software: BS, JD, AEG; Writing - Original Draft Preparation: BS; Writing - Review & Editing: BS, JD, KE, AEG

